# A flexible and efficient microfluidics platform for the characterization and isolation of novel bacteriophages

**DOI:** 10.1101/2022.09.13.507764

**Authors:** Adam Sidi Mabrouk, Véronique Ongenae, Dennis Claessen, Susanne Brenzinger, Ariane Briegel

## Abstract

Bacteriophages are viruses that infect bacteria. This property makes them highly suitable for varied uses in industry or in the development of the treatment of bacterial infections. However, the conventional methods that are used to isolate and analyze these bacteriophages from the environment are generally cumbersome and time-consuming. Here, we adapted a high-throughput microfluidic setup for long-term analysis of bacteriophage-bacteria interaction and demonstrate isolation of phages from environmental samples.

## INTRODUCTION

Bacteriophages are viruses that are highly specialized to infect a narrow spectrum of bacterial species. They are thus a major driving force of bacterial evolution and the structure of environmental bacterial communities (1, 2).

Phages recognize their host via specific receptors such as sugars or proteins exposed on the outside of the cell. Once they are irreversibly bound to the cells’ surface, the phages eject their own genome, consisting of DNA or RNA, into the host cell and reprogram the cell to either produce new progeny (lytic lifecycle) or insert their genome into the host chromosome (lysogenic lifecycle) (3). The high level of host specificity paired with their ability to efficiently decimate their host species has increased interest in bacteriophages as a potential treatment for bacterial contamination in industrial settings (4) or infections from antibiotic resistant bacteria in clinical settings (3, 5–7).

Especially in light of their potential practical application, it is important to isolate and characterize new phages to increase the diversity of species-specific phages at our disposal. However, isolation of phages from the environment and studying their characteristics such as host spectrum and speed of eradicating a population, is time-consuming as traditional methods are hard to translate in a high-throughput manner (8). A proven method within the bacteriophage literature advises to concentrate the environmental sample first by ways of tangential flow filtration to increase the concentration of bacteriophages in the sample (9). At this point, the sample may contain a multitude of different bacteriophages and needs to be enriched for the phages targeting the host of interest. This is typically performed by adding the sample to a liquid culture of the presumed host and, after an incubation period of 24-48 hours, taking the cleared lysate to perform a plaque forming units (PFU) assay on double agar overlay plates to isolate the phages (10). These steps present several problems: i) The nature of the preprocessing steps, phage enrichment and PFU assay result in an isolation protocol that is both time-consuming but also spatially inefficient, due to the lab space that performing these assays with a multitude of samples would occupy. ii) The enrichment during co-culture selects for lytic phages with large burst sizes, fast replication cycles, and/or phages that do not promote establishment of a resistant host within the cultivation time. Phages with slower replication cycles or development of resistant hosts may be missed due to absent clearing of the culture. iii) Furthermore, the intensive filtration steps likely biases against larger sized phages, such as jumbo phages, due to the small pore size of the filters (12).

In addition to the problems described during isolation of phages, the long-term microscopic observation of their interaction with their host is challenging. On microscope slides, bacteria are prone to drying, starvation and lack of oxygen while the movement of phages in higher-percentage agarose typically used to fix motile bacteria may be constricted. Classical microfluidic setups that allow longer observation windows are expensive and require external supplies such as pumps that may not be available to many labs.

In an initial application, we have previously tested the application of a versatile plate-based platform to study bacteria-phage interactions (13). Here, we thoroughly test, expand and describe this setup to allow the screening of bacteriophage activity in a spatially efficient and high-throughput manner without the need of external pumps or tubing. Furthermore, it allows for the direct investigation of a wide range of bacterial behavior in response to phage exposure. For example, we can directly observe bacterial cell wall-shedding upon phage exposure. In this study, we tested this setup on various known bacteria and phage combinations before using the system to isolate a novel phage from an environmental sample.

## RESULTS

### Bacteriophage activity is detectable with E. coli at different titers and with different phages

To develop the desired setup, we aimed to use a commercially available high-throughput microfluidics platform. The Mimetas Organoplates®, developed to be used with cell cultures and organoids, offers various advantages: Each plate contain 96 chips, thus, thus up to 96 bacteria-phage pairs can be studied in parallel. Each chip consists of two channels that merge into one interaction space with only a narrow curb (phaseguide) between the channels (Fig. 1) (14, 15). One channel can be filled with a matrix with embedded bacteria, forming a meniscus of matrix along the phaseguide. The second channel can be filled with liquid samples that are directly adjacent to this matrix meniscus. Thus, the matrix and liquid are not separated by a membrane or physical barrier and nutrients or particles such as phages can diffuse into the matrix while the host bacteria are fixed. This membrane-free separation due to the phaseguide is essential for a more efficient downstream processing of the enriched phage sample (14). The thin glass at the bottom of the plates allows for high quality imaging or scanning using plate readers which facilitates the continuous observation of any kind of bacteriophage related activity. As these plates are designed for eukaryotic cell culturing, it was vital to establish whether both bacteria and phages were able to propagate in this system.

**Figure. 1.**
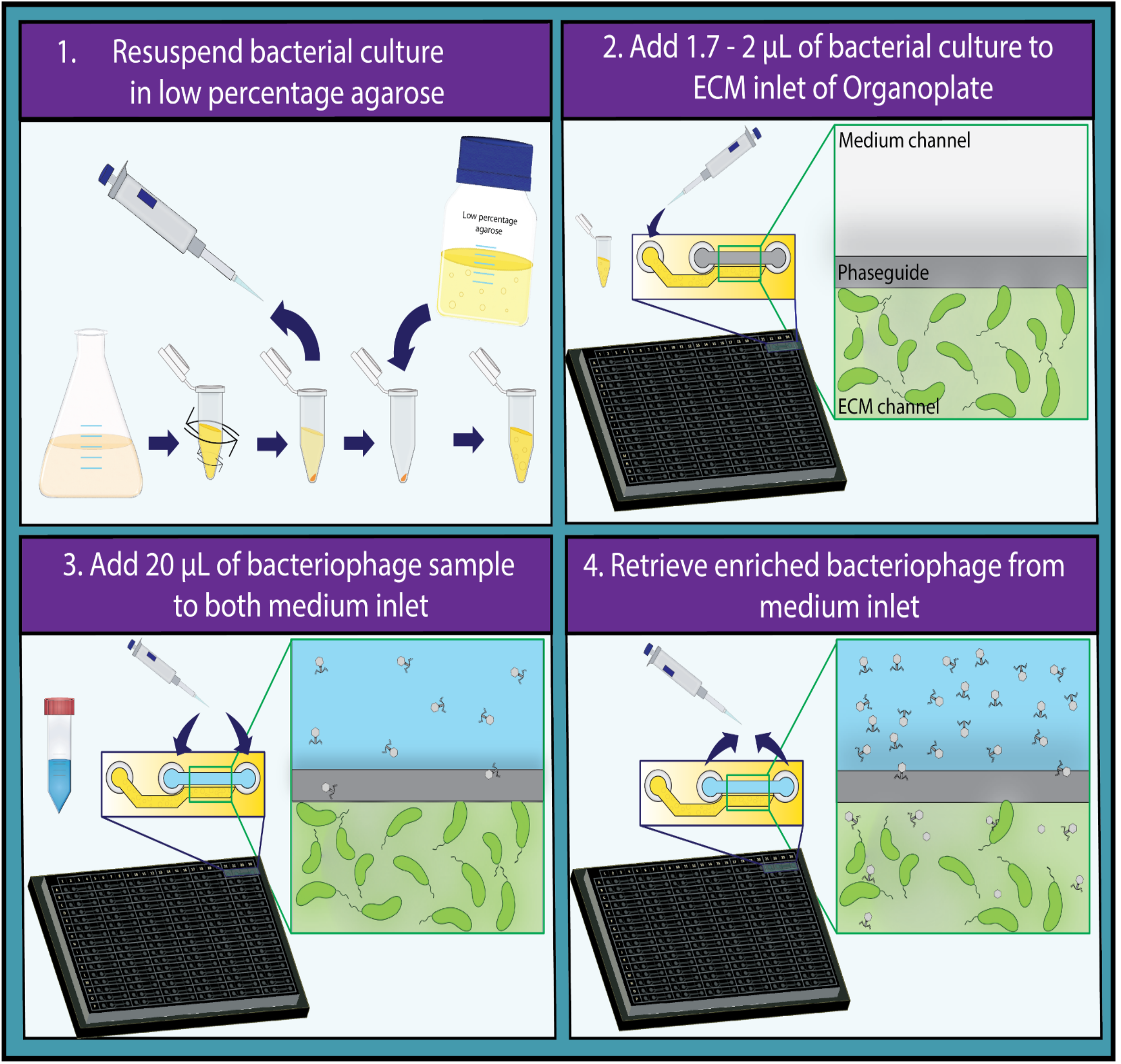
Proposed high-throughput method for the detection and characterization of bacteriophages. Protocol for the use of the high-throughput Organoplate® for the detection of bacteriophage activity divided in four steps. 1. Grow the bacterial culture of interest to exponential phase and resuspend it in growth medium containing 0.4% agarose. 2. Pipette 1.7 – 2 µL of the resuspended bacterial culture in the ECM inlet and let it solidify for 10 minutes. 3. Add 20 µL of phage containing sample to both medium inlet of the plate and rock the plate for 5 minutes. 4. After imaging the plate over a desired period, retrieve the enriched bacteriophage sample from the channels that showcased bacteriophage activity.

To confirm the suitability of the selected platform, we used well established bacteria and phage pairs to ensure that this system was indeed able to facilitate these interactions. Initial tests were done in triplicates with the T4 phage (100 - 0.1 MOI) and an *E. coli* producing sfGFP. Observations were carried out for up to 70 hours on an automated microscope. Compared to the control chips containing only buffer in the liquid channel, the phage treated samples lost fluorescence within ∼20 hours while fluorescence of the control increased continuously (Fig. 2A). This shows that both the embedded bacteria as well as the phages can propagate in the chips. We then further tested the sensitivity of our setup by treating the *E. coli* with a 10-fold dilutions series of T4 phages (10^5^-10^0^ phages per channel) (Fig. S1). We observe a slight overall loss of fluorescence in all conditions over time, both control and phage treated samples. This is possibly caused by decreasing nutrient and oxygen concentrations and/or accumulation of toxic degradation products since it is a closed system. However, this loss of fluorescence did not affect the interpretability of the experiments. Between 10^5^ and 10^3^ bacteriophages, lysis of *E. coli* by T4 was clearly detectable (Fig. S1). Between 10^2^ to 10^0^ bacteriophages a slight difference was only observed after 60 hours of exposure. These results confirm that the system is suitable to test bacteriophage activity and is sensitive enough to detect differences in bacteriophage titer. Next, the plates are tested on a plate reader. Here, a decrease was also observed by automated monitoring of the channels (Fig. S2), further showcasing potential for automation of the setup. Finally, we tested the setup using different known lytic phages of the *E. coli* host, namely T3 and T7 (Fig. 2B). Similarly, these bacteriophages also result in a decrease in fluorescence of the bacteria over time compared to the control, which demonstrates that the system is functional with a variety of lytic phages for a single host.

**Figure 2.**
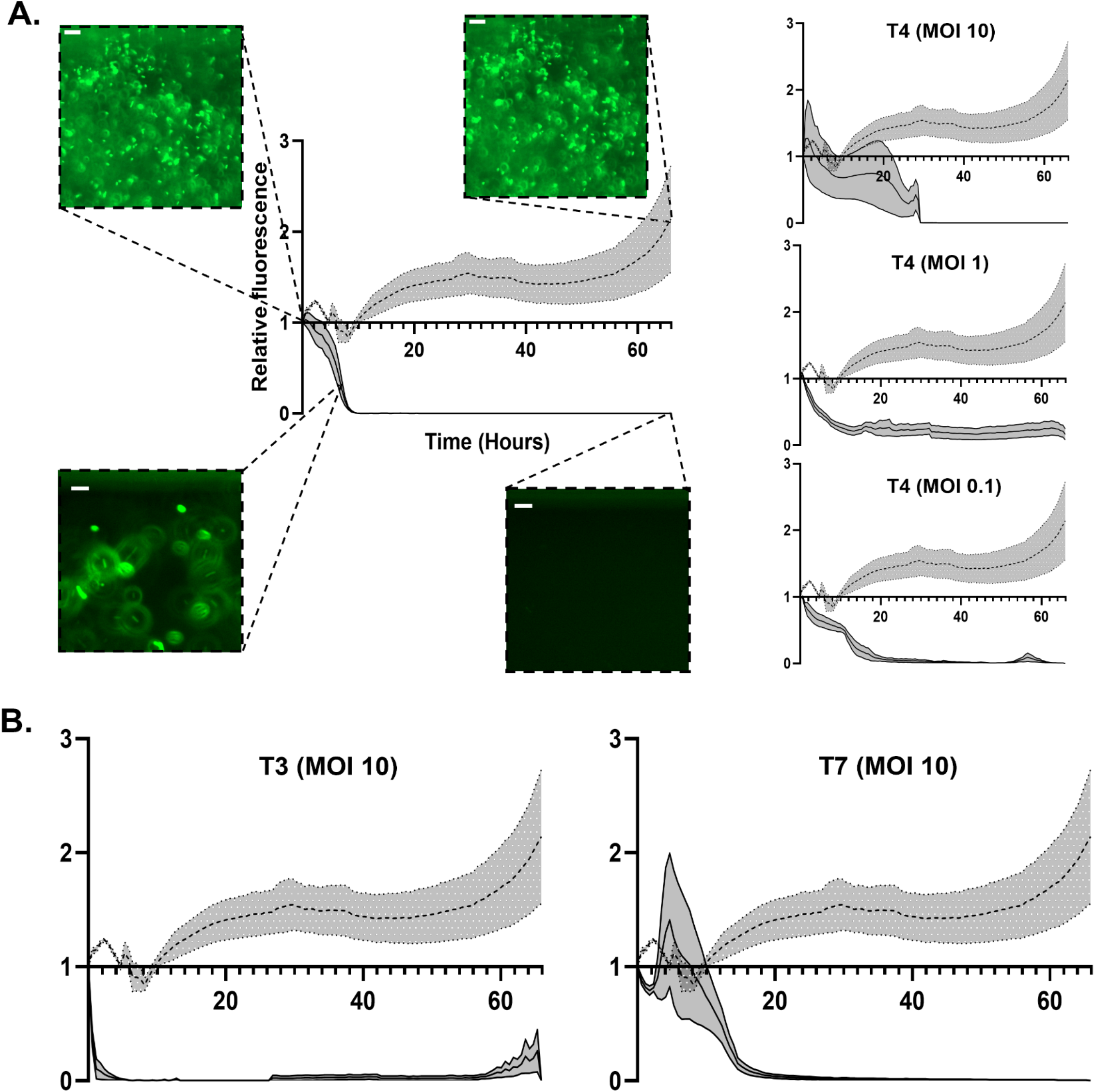
Relative fluorescence measurement of *E. coli* exposed by different phages at different MOI’s. (A) Various graphs depicting the relative fluorescent signal of GFP labeled *E. coli* over a period of 66 hours. The dotted line represents the control sample treated by phage buffer, while the solid line represents the sample that is treated by T4 phages. Different phages and titers are depicted in each graph. Grey zones represent standard mean of error (SEM). Representative images of *E. coli* are depicted at time points 0, 7 and 66 hours for both the control sample and T4 bacteriophage at a MOI of Scale bars represent 15 µm. (B) Graphs depicting the relative fluorescent signal of GFP labeled *E. coli* over a period of 66 hours. The dotted line represents the sample treated by phage buffer, while the solid line represents the sample that is treated by either T3 or T7 phages. Grey zones represent standard mean of error (SEM).

### Bacteriophage activity can be detected for other organisms and in other detectable metrics

To determine if the same setup can be used with other species, we tested a pathogenic diderm and a monoderm host, *Vibrio cholerae* and *Bacillus subtilis* respectively, with their respective phages ICP1 and φ29 (16, 17, 18). Transgenic fluorescently tagged *V. cholerae* N16961 exposed to control buffer samples displayed slower growth compared to *E. coli* and an increase in fluorescence is only visible from 20 hours onwards (Fig. 3A, S3A). Despite this slow growth, there still is a stark difference between the control and the phage-treated sample, as fluorescence disappears very rapidly after the addition of phages (Fig. 3A, S3B). While addition of the phage did not lead to lysis of the cells, another interesting phenomenon was visible here. Both in the control channel and the phage channel, all cells changed from their normal rod-shape to a spherical morphology. While this happened in both conditions eventually in the presence of the φ29 bacteriophage it happened after only 5 hours in contrast to 50 hours in the control setup (Fig. 3B-C). This resembles our previously reported observations which described phage induced escape mechanism that allows the bacterial cell to shed its cell wall and thereby evade phage predation (13, 19). By exposing these spherical cells to water, we observed that the majority of cells were lysed which indicates that these are likely not spores but indeed cell wall-deficient cells as described in Ongenae et al. (13, 19) (Fig. S4). In this case, this system was able to clearly detect whether *B. subtilis* is predated by phages as the morphological transition occurred 45 hours earlier than in the control channel. In addition, our plate-based setup can capture responses to phages beyond mere lysis of cells which would be difficult to detect with conventional methods.

**Figure 3.**
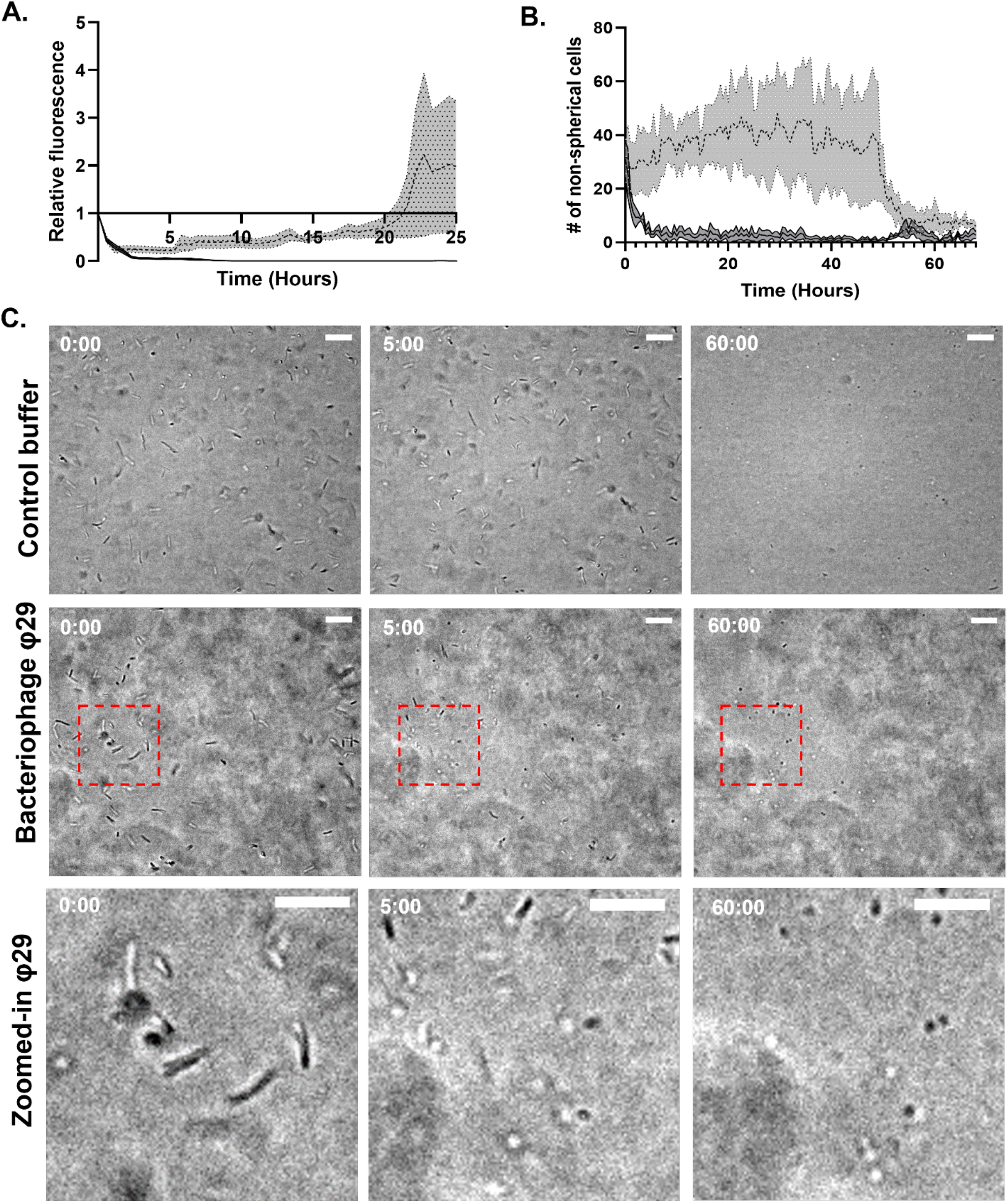
Detection of bacteriophage activity in *V. cholerae* and *B. subtilis*. Fluorescent signal relative to the first measurement of GFP labelled *V. cholerae* over a period of 25 hours. Dotted line represents control sample that is exposed to phage buffer. Solid line represents *V. cholerae* that is exposed to ICP bacteriophages at a MOI of 10. Grey zones represent standard mean of error (SEM). (B) Graph depicting the amount of non-spherical *B. subtilis* cells over a period of 66 hours. The dotted line represents the sample treated by phage buffer, while the solid line represents the sample that is treated by the φ29 bacteriophage at a MOI of 10. Grey zones represent standard mean of error (SEM). (C) Images at specific time points depicting the change of *B. subtilis* cells from rod-shaped to spherical. The control, phage-treated and a zoomed-in version of the phage-treated condition are depicted respectively from top to bottom. Time in hours is depicted in the top left corner. Scale bars represent 15 µm. Note: Image shift between timepoints 0:00 and 5:00 sometimes occurs during data collection in the automatic setup.

### Detection of a novel E. coli bacteriophage from a sewage treatment plant sample using the high-throughput microfluidic setup

With the setup being established to detect bacteriophage activity for multiple different bacteria, it could now be applied for the detection of bacteriophages in environmental samples specific to these bacteria (see workflow in Fig. 1). For this purpose, we collected a water sample in close proximity to a sewage treatment plant, since these sites are described as areas that contain plenty of bacteriophages due to the wide diversity and abundance of bacteria that are present in the water (20). The previously tested organisms, namely *E. coli, V. cholerae* and *B. subtilis*, were exposed to this sewage water in the high-throughput microfluidics plates. If the phages that are present in the water are specific to either *E. coli, V. cholerae* or *B. subtilis*, we expect a similar response to that of the previously performed experiments with the known species-specific phages. The addition of the water sample did not seem to affect the latter two organisms but did clearly result in a decrease in fluorescence of *E. coli* (Fig. 4, S5). Predation of this phage on *E. coli* happened rapidly, showing a significant reduction as early as 9 hours after the addition of the water (Fig. 4). Subsequently, we observed an increase of fluorescence over time, accompanied with possibly resistant bacteria occurring in the channels (Fig. 4).

**Figure 4.**
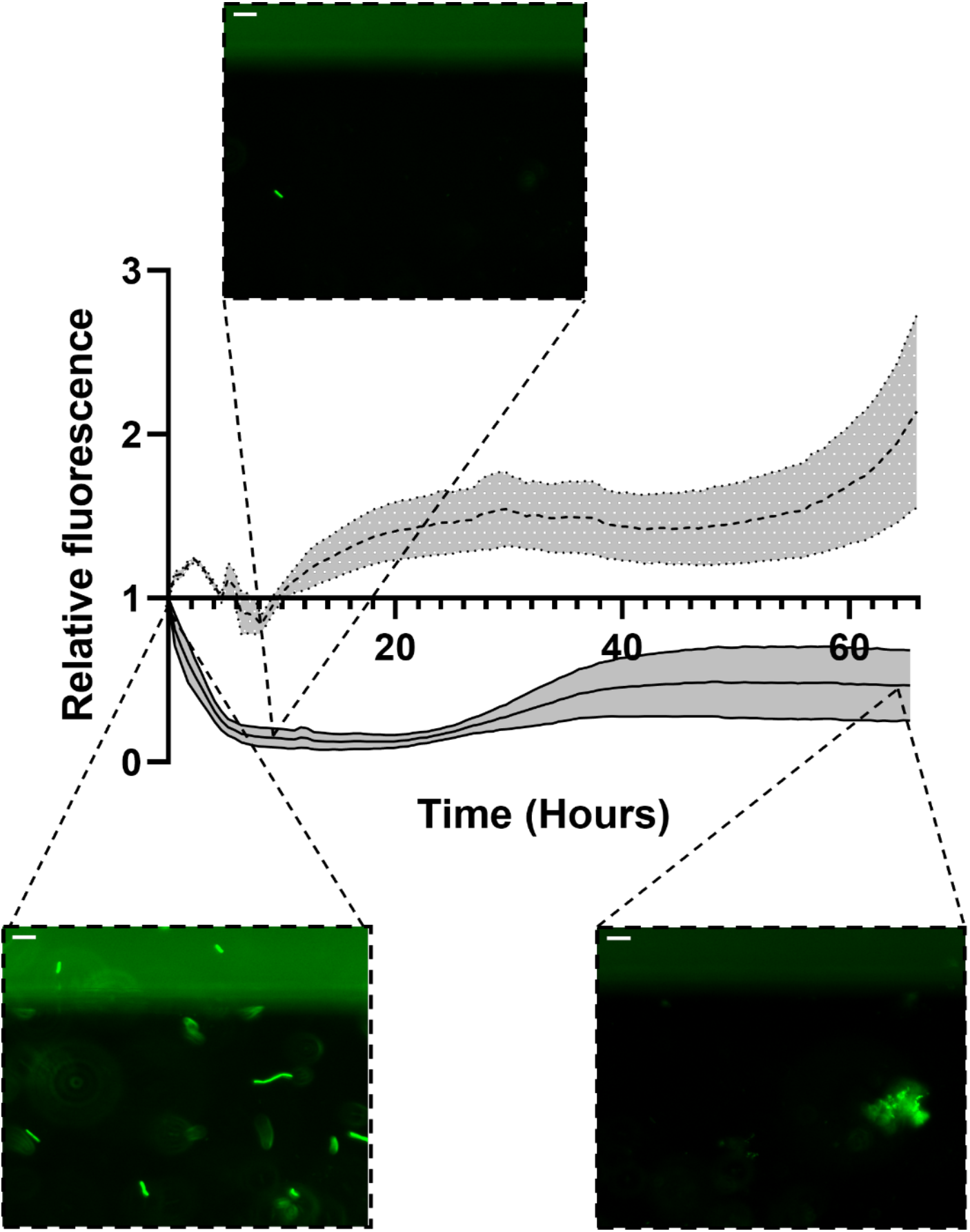
Clearance of fluorescent *E. coli* exposed to phages from an environmental water sample. Fluorescent signal relative to the first measurement of GFP labelled *E. coli* over a period of 66 hours. Dotted line represents control sample that is exposed to phage buffer. Solid line represents *E. coli* that is exposed to the environmental water sample. Grey zones represent standard mean of error (SEM). Representative images of *E. coli* are depicted at time points 0, 9 and 66 hours for the environmental water sample. Scale bars represent 15 µm.

The total remaining volume of the water sample (∼40 µL) was recovered from the specific channels that resulted in a decrease of bacteria over time and a subsequent plaque assay was performed. This was done to both confirm the presence of the phages in the water and to subsequently isolate the enriched coliphages.

After isolation of the phages from the water, genome isolation and subsequent sequencing was performed to confirm the identity of the phage. Sequencing showed that this phage is an unidentified species of coliphage belonging to the podovirus family. The closest relative is another recently identified phage called PTXU04 (21). The genome of the phage, which was named Pondi, has a size of 61061 bp and 86 open reading frames (ORFs) could be identified (Fig. 5B). To confirm the morphology of the phage, we imaged the sample using cryo-electron tomography (cryo-ET). The imaging revealed a typical podovirus-like morphology when the capsid head was full of DNA (Fig. 5A). The empty-head phages (those that had ejected their DNA) further revealed that the tail is retractable, since its length varied greatly compared to the phages with a full capsid head (26.55 ± 1.57 nm compared to 51 ± 3.89 nm). Furthermore, flexible tail fibers were identified in these images. The successful isolation of this new phage demonstrates that this setup is indeed capable of identifying novel phages from environmental samples without the need to amplify your phage sample first.

**Figure 5.**
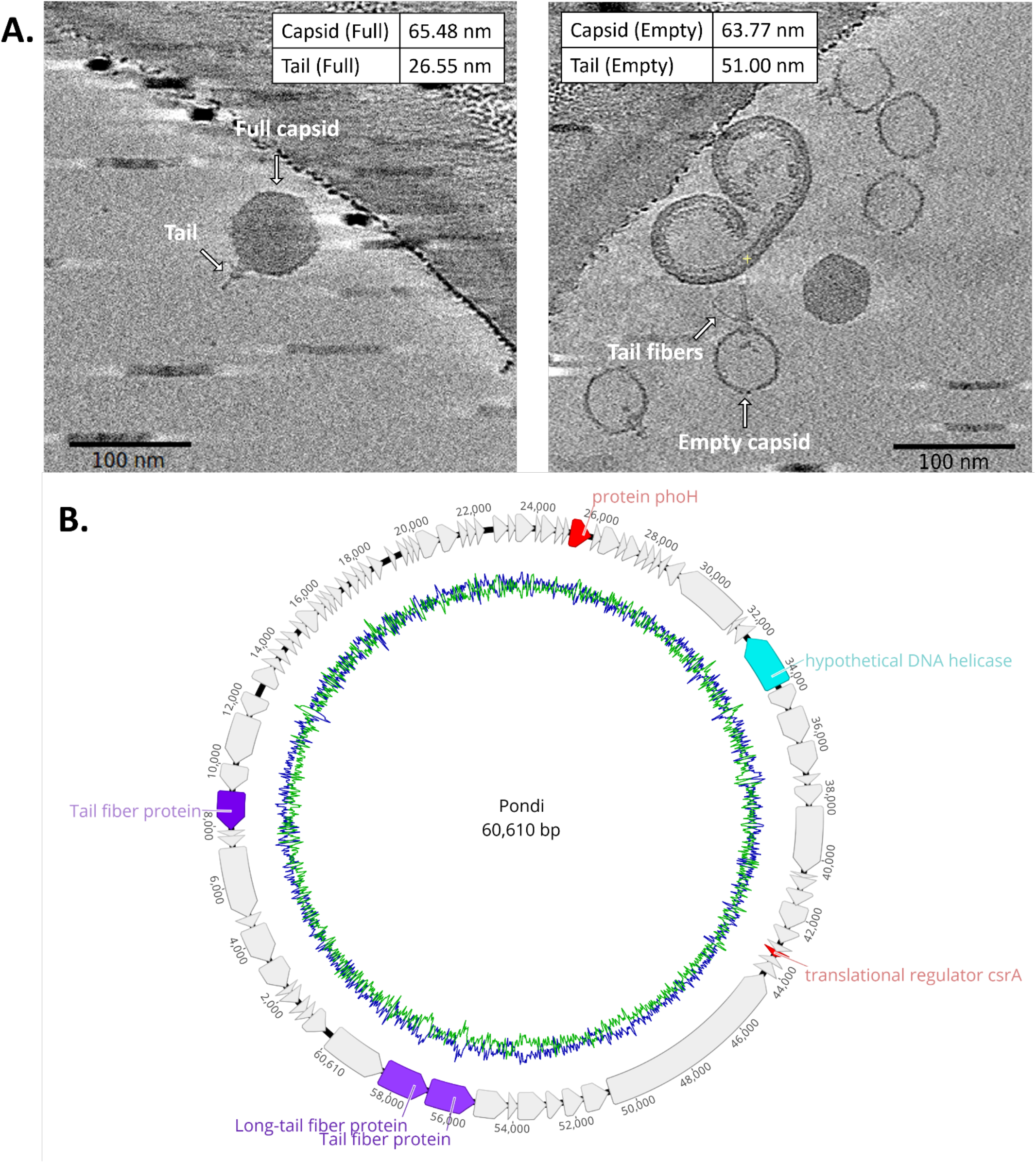
Characterization of the novel bacteriophage Pondi. (A) Cryo-ET images of the newly identified *E. coli* phage. Measurements of the dimensions for full (N = 5) and empty (N = 9) capsid phages in both images. Scale bars represent 100 nm. (B) Genome annotation of Pondi. Each individual ORF is color-coded. Grey, purple, red and teal represent hypothetical proteins, structural proteins, bacterial proteins and DNA regulatory proteins, respectively. G+C content is represented in blue, while A+T content is represented in green.

## DISCUSSION

The awareness of the potential use of bacteriophages for various applications in industry and medicine is steadily increasing. However, practical applications will greatly rely on the availability of suitable phages. At present, the content of phage libraries around the world are still limited, and the need for new specific phages for a large array of bacterial hosts will only grow in the near future (22, 23). Here, we aimed to develop a screening platform with increased efficiency and ease of phage sample characterization and isolation from environmental samples compared to traditional methods.

The Mimetas Organoplates® was chosen over simple 96-well dish plates for two reasons. First, the quality of imaging that is possible with the Mimetas Organoplates® is vastly superior compared to what is achievable with a 96-well plate. The thin glass bottom, fixation of motile bacteria in a matrix and barrier-free separation of bacteria and phages by the phaseguide allows for improved long-term imaging that can capture small details such as a change in bacterial cell shape. This is illustrated by the observation of the formation of cell wall-deficient cells in response to phage predation (13). Likely, this behavior went unnoticed previously due to the limited imaging quality possible with a 96-well plate setup.

We were able to investigate the phage-induced shedding of the cell wall in several species by observing the transition from rod-shaped to spherical cells. However, we observed that *B. subtilis* cells also became spherical in the absence of phages (Fig. 3C). This is likely due to limited oxygen or nutrient starvation and/or accumulation of waste products (24), as it only happens after a prolonged amount of time without the addition of any new nutrients. Alternatively, these spherical bodies that form under prolonged growth in the Mimetas setup in the absence of phages could be due to the formation of spores. *B. subtilis* is capable of producing spores in conditions of environmental stress (25). However, the striking similarity in size and appearance to the wall-less cells that form under phage attack make this interpretation possible but less likely. In either case, the setup allows not only for the isolation of new phages, but simultaneously allows for the study of the host response.

Secondly, the membrane-free separation of phages and bacteria in the system has multiple advantages. Downstream processing of the environmental sample after enrichment does not require the separation of phages and bacteria by filtration or precipitation techniques, which has been implied to alter the viability of the phages (26). Additionally, this also paves the way for high-throughput processing.

Various other high-throughput methods have been described for the isolation of bacteriophages. One other method uses 96-well plates and direct plaque sequencing (DPS) methods to achieve a faster throughput (27). Here, all 96 samples are analyzed indiscriminately of any kind of phage activity, with the sequencing results acting as the confirmation of the presence of any bacteriophages. There are a couple of downsides to this method compared to ours. With 96 well-based systems, it is not possible to monitor the effect on the host bacteria overtime, while we can accurately judge which channels exhibit bacterial responses that are indicative of phage activity. This reduces the amount of downstream work necessary as not all the samples are selected for further processing, just the ones that possibly contain bacteriophages. Furthermore, 96 well-based systems require a subsequent plaque assay that is performed on one double agar overlay plate which limits the setup to only use one specific host. In contrast, our setup allows for using multiple hosts (with similar growth conditions) simultaneously. If several host cells are tagged with different fluorophores, they could also be used within the same chip, allowing further upscaling of our setup. Overall, our setup is an effective way to reduce the time and space necessary for large scale phage combination experiments.

Finally, while we have tested the system using known lytic host-phage systems as well as the isolation of the new lytic phage Pondi from an environmental water source, it can also be used to dynamically study other stressors such as drugs, either in combination or separately. As the liquid channels allow fast and easy access to the micro-organisms of interest, it is also possible to administer the drugs at later time points during an experiment. At present, we have not tested this setup using lysogenic phages, since they would be difficult to detect as it would not always lead to a visible decrease in bacterial cells. In future experiments, we aim to use lysogenic host-phage systems and treat them with mitomycin C. This antibiotic can be administered to bacteria carrying a lysogenic phage, which would result in the release of prophages and subsequent lysis of the infected cells (28), making this platform also suitable for lysogenic phages.

In summary, we showcase a new setup for phage isolation and studying the host response at high resolution using an adapted high-throughput microfluidic system. This flexible and adaptable system will ameliorate the current tools that are available for the detection of bacteriophages. With the current untapped potential for bacteriophages, both in industry and in the clinics, the prospect of possibly finding novel bacteriophages with this system becomes even more exciting.

## MATERIALS AND METHODS

### Growth conditions and strains

All strains used in this study are listed in Table 1. Bacterial strains were cultivated under their respective standard laboratory conditions in lysogeny broth (LB) with shaking. All strains were preserved in frozen stocks and streaked on fresh LB plates for experiments. Colonies from these plates were used for inoculation of liquid cultures. These cultures were grown overnight and used for reinoculation of fresh liquid cultures to an OD_600_ of 0.05 unless mentioned otherwise. Induction of the pBTOK-sfGFP plasmid was done by the addition of anhydrotetracycline (20 ng/mL) 45 minutes before the start of the experiment. All bacteriophages were stored at 4°C in phage storage buffer (100 mM NaCl, 10 mM MgSO_4_, 10 mM Tris-HCl and 1 mM EDTA). Propagation of the phages was performed on their respective host in LB media.

**Table 1:**
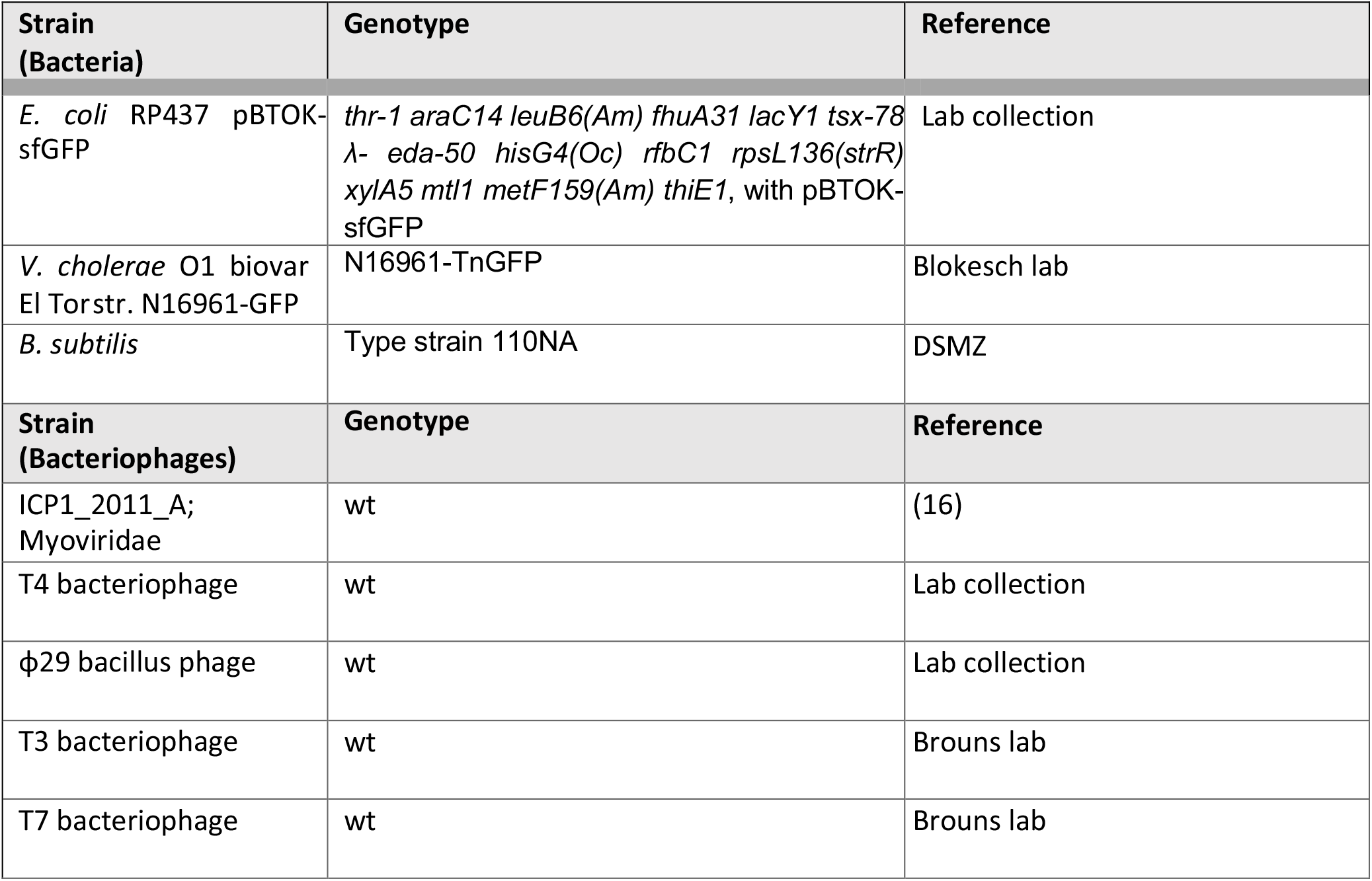
Bacterial strains and bacteriophages used in this study.

| **Strain**<br>**(Bacteria)** | **Genotype** | **Reference** |
| --- | --- | --- |
| *E. coli* RP437 pBTOK-sfGFP | *thr-1 araC14 leuB6(Am) fhuA31 lacY1 tsx-78*<br>λ- *eda-50 hisG4(Oc) rfbC1 rpsL136(strR)*<br>*xylA5 mtl1 metF159(Am) thiE1*, with pBTOK-sfGFP | Lab collection |
| *V. cholerae* O1 biovar El Torstr. N16961-GFP | N16961-TnGFP | Blokesch lab |
| *B. subtilis* | Type strain 110NA | DSMZ |
| **Strain**<br>**(Bacteriophages)** | **Genotype** | **Reference** |
| ICP1_2011_A;<br>Myoviridae | wt | (16) |
| T4 bacteriophage | wt | Lab collection |
| ϕ29 bacillus phage | wt | Lab collection |
| T3 bacteriophage | wt | Brouns lab |
| T7 bacteriophage | wt | Brouns lab |

### Plaque assay

A fresh overnight culture of the respective host bacteria was diluted to an OD_600_ of 0.05 in 10 ml LB. After reaching an OD_600_=0.3-0.6, 1 ml of the culture was combined with 5 mM CaCl_2_ and 50 ml soft agar (0.3% w/v agar in LB) that was cooled down to 37°C. The culture was mixed and 12.5 ml was pipetted onto prewarmed LB plates. The culture was solidified at room temperature for 30 minutes. A gradient of phage stock was made from 10^7^ to 10^11^ PFU/ml and 3 μl of each concentration was carefully pipetted on the plate. The plates were incubated overnight at room temperature after which the plaque counts were determined.

### High titer phage stock preparation

A 50 ml LB culture of the respective host bacteria (e.g. *E. coli, V. cholerae* or *B. subtilis*) was grown to an OD_600_ of 0.3-0.6 followed by the addition of the respective bacteriophage (e.g. T4, ICP1, φ29) at a concentration of 10^9^ PFU/mL. After a 6-hour incubation period, the cells were centrifuged for 30 minutes at 5000 rpm (rotations per minute) at 4°C. The supernatant was filtered through a 0.45 μm filter and incubated with phage precipitation solution (4% PEG 8000, 0.5 M NaCl) at 4°C overnight. The precipitated phages were collected by centrifugation at 3000 g for 1 hour at 4°C. Collected phages were resuspended in phage storage buffer (100mM NaCl, 10mM MgSO4, 10mM Tris-HCl, 1mM EDTA). The phage titer was determined by plaque assay of a serial dilution as previously described.

### Mimetas Organoplate® protocol

An overnight liquid culture of the bacteria of interest was grown at the appropriate conditions. The OD of the overnight culture was measured and diluted to an OD_600_ of 0.1. Subsequently, the host culture was grown until it reached an of OD_600_ of 0.3-0.5. Meanwhile, 0.4% agarose (Molecular Biology Grade Agarose, Eurogentec, Belgium) was heated up in LB medium in the microwave and kept at 45°C to prevent solidification. When the host culture had reached an OD_600_ of 0.3-0.5, 1000 μL was spun down at 5000 rpm for 10 minutes. The supernatant was removed and the pellet was resuspended in 0.4% body temperature soft agarose to a final theoretical OD_600_ of 0.3. CaCL_2_ (50 mM) was added to the resuspended culture to aid phage adsorption. 1.7 - 2 μL of resuspended culture was pipetted in the ECM inlet of the Mimetas 2-lane Organoplate® (Fig. S1). This was repeated until enough lanes contained the bacterial culture of interest. Afterwards the pipetted agarose was left to solidify for 10 minutes. After the agarose matrix had solidified, 20 μL of either bacteriophage, control or environmental sample was added to both medium inlets. Each condition was performed in triplicates, meaning in three separate chips. The plate was gently rocked for around 5 minutes to ensure distribution of the bacteriophage. The plate could now be used to screen for clearance. For this purpose, either a plate reader or microscope to continuously measure clearance over time was used.

### Fluorescence microscopy imaging

Imaging was performed with the Lionheart FX automated microscope (Biotek, Winooski, USA). Depending on the strain, imaging was performed either with or without fluorescence readout (Ex: 469 nm, Em: 525 nm). Below the phaseguide, multiple areas of the gel inlet were imaged (Fig. 1). The magnification was set to 20x and the automated laser autofocus function was used to determine the height at which imaging happened. A reference image was acquired to ensure the microscope was adjusted for the desired area on the Mimetas Organoplate®. A montage of five different areas within the channels of the plate was made to cover a wider imaging area. Imaging was done overnight, varying between 24-72 hours of total imaging time. The imaging chamber was kept at 30°C to ensure optimal growth conditions for the bacteria. Clearance of bacteria was measured over time by means of loss of fluorescence or by the number of spherical cells. Quantification was performed in Fiji by means of pixel threshold counting for the loss of fluorescence The “analyze particles” function in Fiji was used to determine the amount of non-spherical cells for the *B. subtilis* cells by only selecting particles with a circularity between 0-0.7. Three different areas of interest were selected of the montage of each chip for the quantification.

### Plate reader

Automated fluorescence measurements was performed with the Spark® multimode microplate reader (Tecan, Männedorf, Switzerland). A Mimetas Organoplate® which contained GFP labelled *E. coli* RP437 and T4 phages at different concentrations were put in the device and imaged every minute for a period of 16 hours. Both OD and fluorescence was measured over time.

### Environmental bacteriophage isolation

Water was sampled around the water treatment plant (coordinates: 52°10’23.5”N 4°29’31.9”E) in a 50 mL falcon tube. The water was filtered through a 0.45 µM filter to get rid of most other micro-organisms in the sample. This filtered sample was then added to in the medium channel of the Mimetas Organoplate®, with the putative host (*E. coli, V. cholerae, B. subtillis*) embedded in 0.4% agarose + LB. The sample was recovered from the channels that showed clearance of the host strain and subsequently used for a plaque assay. Plaques were swabbed and streaked out on a bacterial lane on the host of interest. From this plate, single plaques were selected and picked into a liquid culture of the host of interest at an of around OD_600_ of 0.5. The cleared lysate was finally used to make a high titer phage stock.

### Cryo-ET

Novel bacteriophage Pondi was isolated and enriched to a concentration of 10^12^ PFU/ml. Quantifoil R2/2 200 mesh carbon grids (Quantifoil Micro Tools, Jena, Germany) were plasma cleaned using the Quorum Q150s glow discharger (Quorum technologies, Lewes, United Kingdoms). Aliquots of 3 μl were added to the plasma cleaned grids and subsequently plunge-frozen in liquid ethane using a Leica EMGP (Leica microsystems, Wetzlar, Germany). Blotting time was set to 1s with the chamber temperature and humidity being at 20°C and 85%, respectively. Samples were transferred in grid boxes (MiTeGen, Ithaca, NY) and stored in liquid nitrogen until use. Images of the grids were collected with a CS-corrected Titan Krios (TFS). A tilt series of each target was collected with a dose symmetric tilt scheme between -54° and 54° with 2° increments and a pixel size of 2.64 Å. The defocus was set between -4 µm to -6 µm with a cumulative dose of 100 e^-^/Å^2^.

### Genomic bacteriophage DNA isolation

Genomic material of novel bacteriophage Pondi was isolated using the Phage DNA Isolation Kit from Norgen (Thorold, Canada) according to the manufacturers protocol. Subsequent whole-genome sequencing, *de novo* assembly and annotation and prediction of ORFs was performed by BaseClear using Illumina NovaSeq PE150 sequencing with an average coverage of 144.05 (Leiden, The Netherlands). The genome sequence of bacteriophage Pondi can be found under accession number: OP136151. The genome was visualized by constructing a circular map with Geneious Prime 2022.2.1.

## Data accessibility

All data are available in the main text and electronic supplementary material. Phage genome sequencing data is available on NCBI (Genbank: OP136151).

## Acknowledgements

We would like to thank Dr. W. Noteborn for assisting with the collection of the cryo-ET data from the NeCEN centre (Leiden, Netherlands) and N.C. Brüchle for help with the use of the platereader. We are also thankful to Dr. M. Blokesch for providing the transgenic *V. cholerae* strain N16961-GFP, Dr. S.J.J. Brouns for providing the T3 and T7 bacteriophages and for Dr. A. Camilli for providing the ICP1 bacteriophage.

---

**Figure S1.**
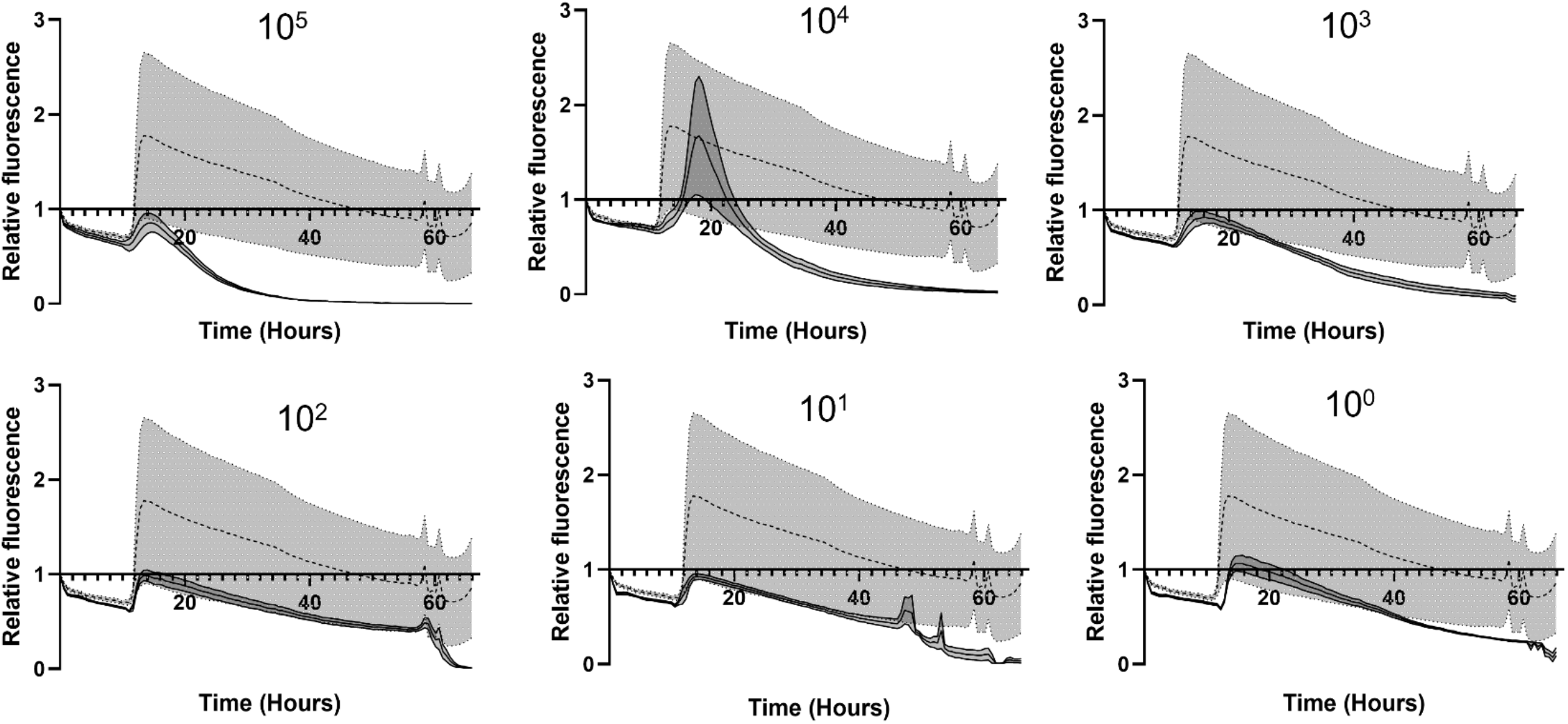
Relative fluorescence measurement of *E. coli* exposed by T4 at the lowest possible concentrations. Various graphs depicting the relative fluorescent signal of GFP labeled *E. coli* over a period of 66 hours. The dotted line represents the control sample treated by phage buffer, while the solid line represents the sample that is treated by T4 phages. The amount of phages used is depicted in each graph. Grey zones represent standard mean of error (SEM).

**Figure S2.**
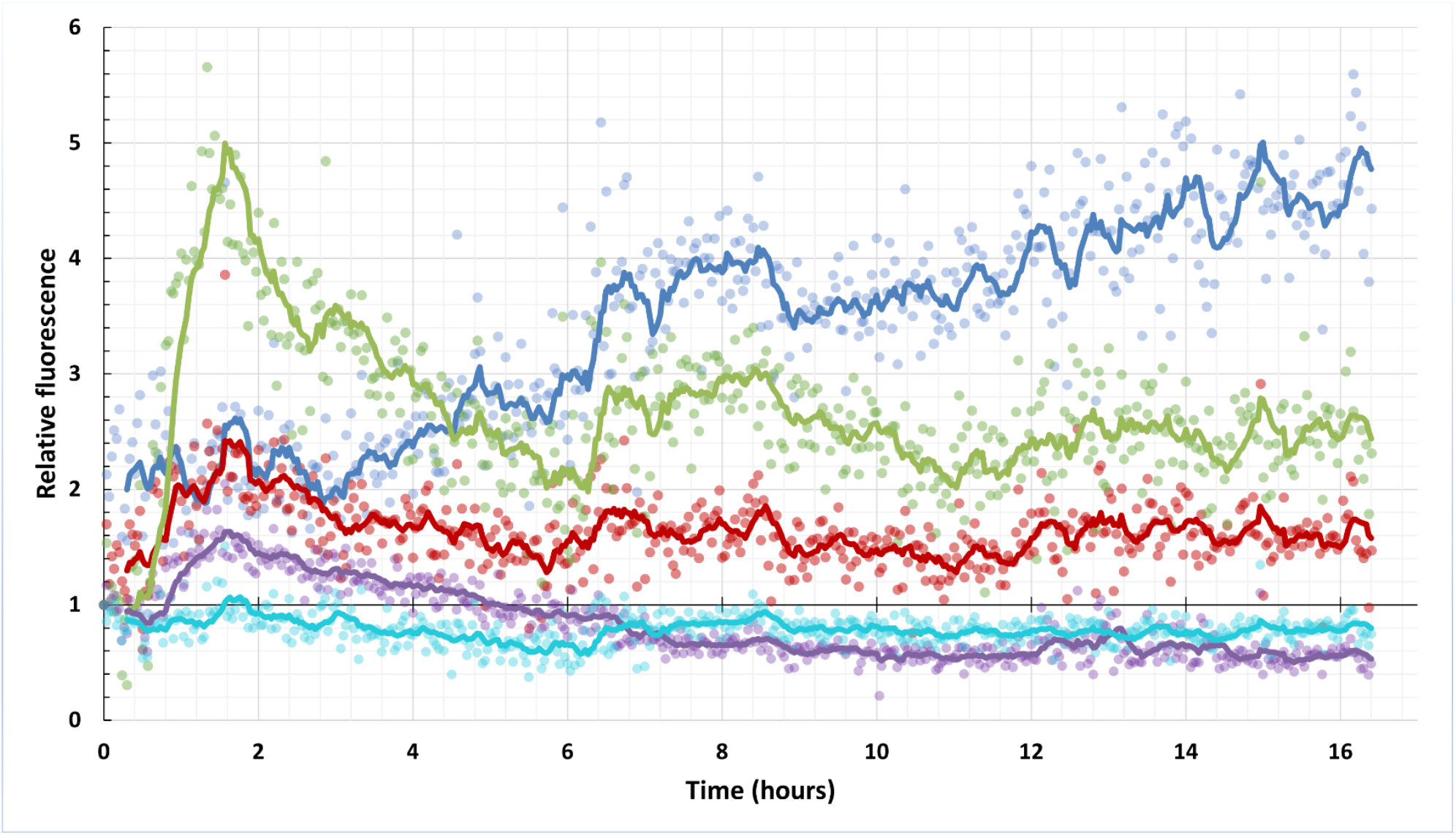
Plate reader data of Relative fluorescence of GFP labeled *E. coli* exposed to T4 at different MOI’s. Graph depicting fluorescent signal relative to the first measurement of GFP labelled *E. coli* over a period of 16 hours. Treatment by T4 at different concentrations and the control are represented by different colors. The control is represented in blue. A MOI of 0.1, 1. 10 and 100 is represented by red, green, purple and teal respectively. The solid line represents the moving average (period: 10) of each respective treatment.

**Figure S3.**
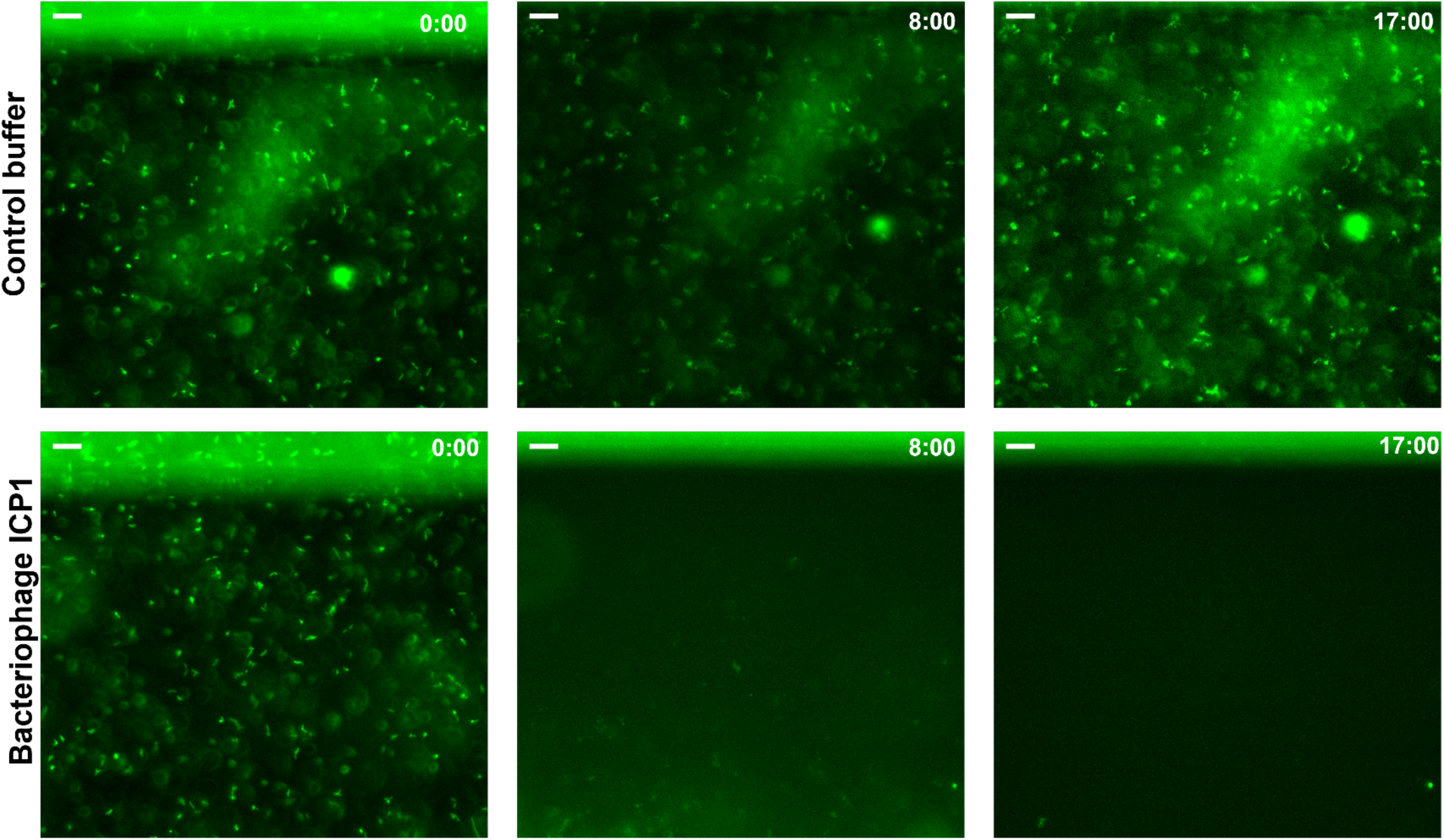
Images of GFP labeled *V. cholerae* exposed to control buffer or ICP1 (MOI: 10) at different timepoints. Images at specific timepoints depicting the effect of ICP1 exposure to *V. cholerae* cells over time. The control (A) and phage treated (B) conditions are depicted respectively from top to bottom. Time in hours is depicted in the top right corner. Scale bar is 15 µm.

**Figure S4.**
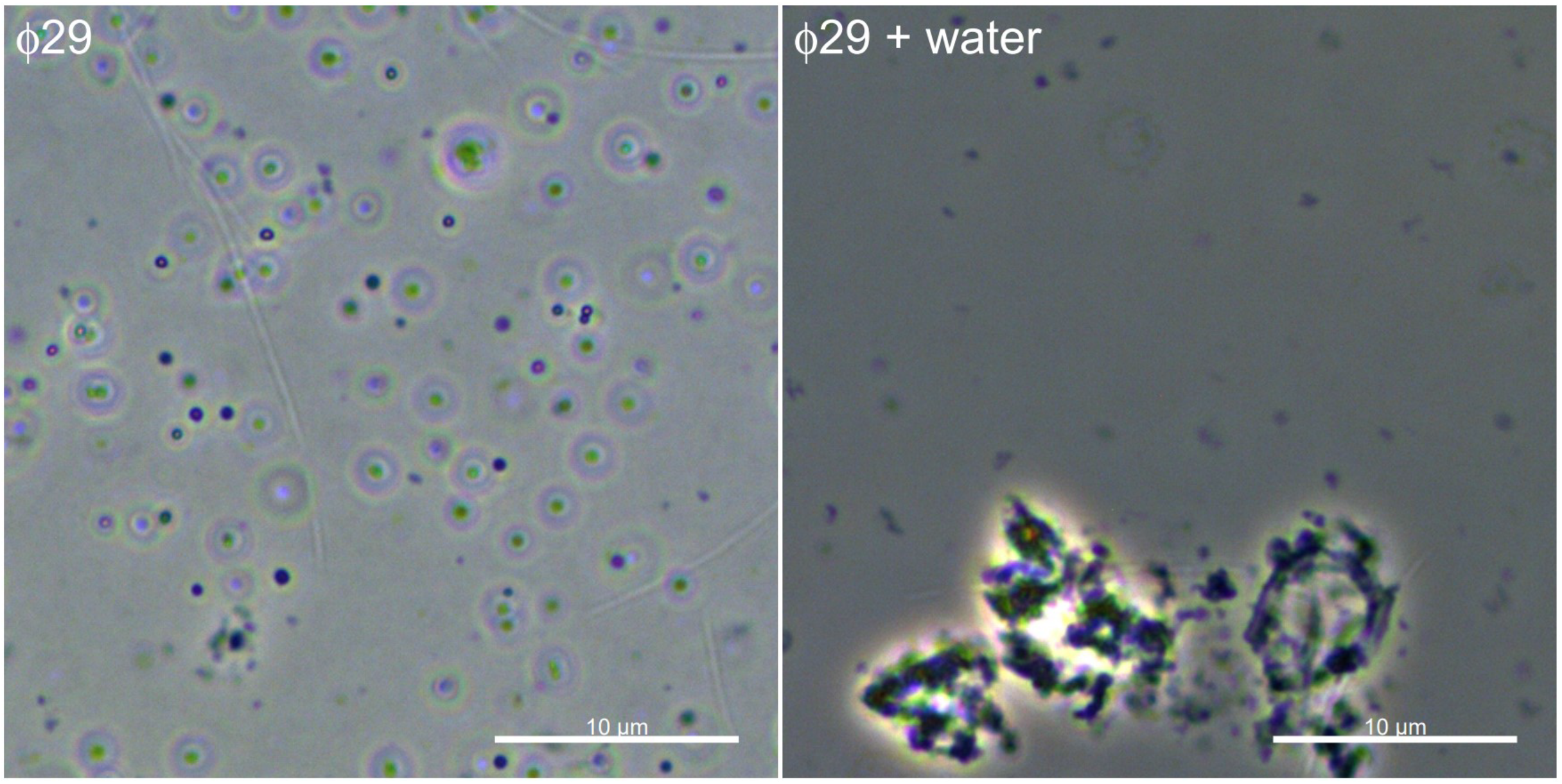
Images of *B. subtilis* cells after exposure to bacteriophage φ29 and to water. Images depicting cell wall-deficient *B. subtilis* cells. These cells can be seen on the left after exposure to bacteriophage φ29. The image on the right showcases the same cells after the addition of water. Scale bar is 10 µm.

**Figure S5.**
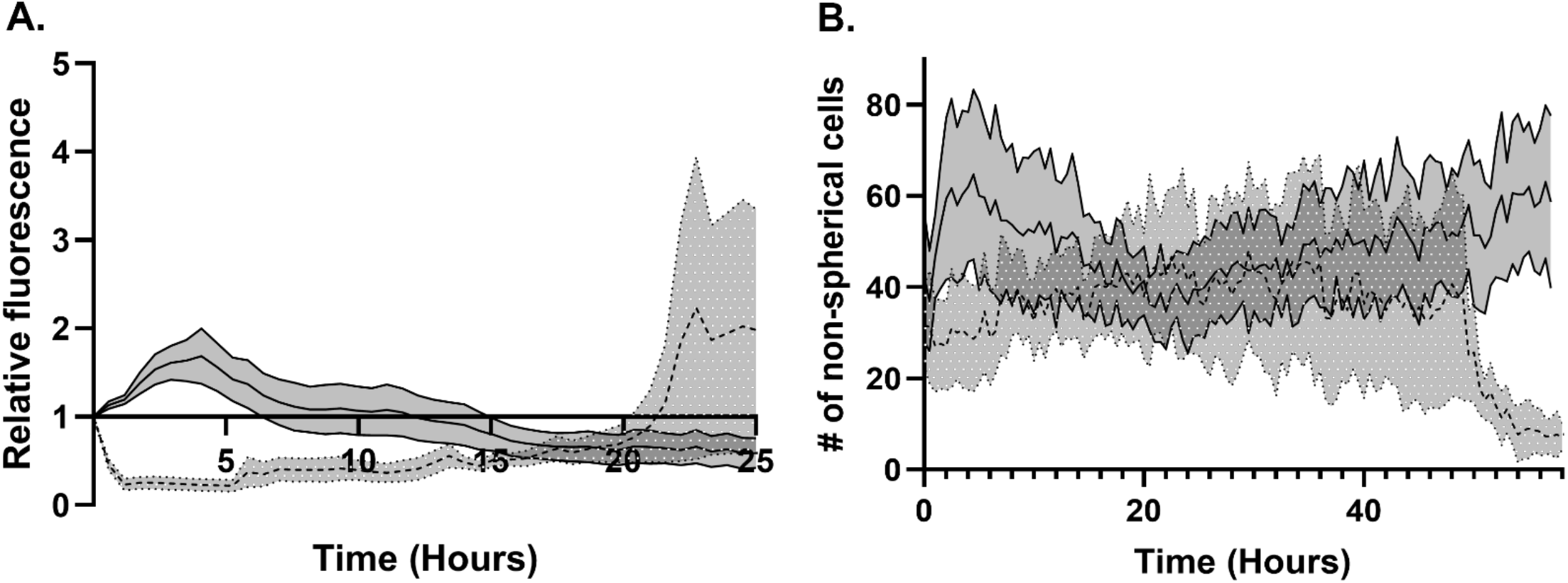
The effect of environmental water sample to *V. cholerae* and *B. subtilis*. (A) Graph depicting fluorescent signal relative to the first measurement of GFP labelled *V. cholerae* over a period of 25 hours. Dotted line represents control sample that is exposed to phage buffer. Solid line represents *V. cholerae* that is exposed to the environmental water sample. Grey zones represent standard mean of error (SEM). (B) Graph depicting the amount of non-spherical *B. subtilis* cells over a period of 66 hours. The dotted line represents the sample treated by phage buffer, while the solid line represents the sample that is treated by the environmental water sample. Grey zones represent standard mean of error (SEM).

